# Effects of mutant and non-mutant strains of *Bacillus subtilis* to produce *ogiri* from *Citrullus vulgaris* seeds on their microbial diversity and physicochemical properties

**DOI:** 10.1101/2024.10.29.620891

**Authors:** Catherine Y. Babatuyi, Victor O Oyetayo, Felix A Akinyosoye

## Abstract

The desire for safe and natural seasoning is increasing as a result of side effects associated with artificial seasonings. *Ogiri*, one of such natural seasonings is a fermented product with different substrates in Africa, especially West Africa. Isolated *Bacillus subtilis* strains from fermented *Citrullus vulgaris* were exposed to different Ultraviolet (UV) irradiations and Sodium Dodecyl Sulphate (SDS) at varying intervals of time to obtain mutant strains. Eight (8) mutant strains of *B. subtilis* that produced high D-ribose metabolites were used for controlled starter-fermentation of *C. vulgaris* for 5 days to produce *ogiri*. The non-mutant (NMS00) and the market (RTE00) served as control samples. Microbial ecology (load and occurrence) and colour characteristics of the samples were determined. The pH increased with the fermentation period from 4.08 ± 0.01 to 9.86 ± 0.01, while the total titratable acidity fluctuated with the fermentation days. The microbial count increased from 1.0 × 10^3^ cfu/g to 5.2 × 10^4^ cfu/g. *B. subtilis* was mainly isolated from *ogiri* samples fermented with mutant strains, while other microorganisms were isolated from samples NMS00 and RTE00. Sample fermented with a mutant strain of *B. subtilis* exposed to inoculating chamber UV light for 90 sec (MIC67) was closer to sample RTE00 in colour measurements. Some of the *ogiri* produced from mutant strains of *B. subtilis* had improved colour characteristics and were not characterized by spoilage or pathogenic microorganisms associated with the market sample. Therefore, could be incorporated into food systems.

**Importance:** Genetic modification of *Bacillus subtilis* strains for the fermentation of *Citrullus vulgaris* seeds to produce a food condiment (*ogiri*), which modified and controlled the kind of microbial load, occurrence and isolations during the fermentation. This prevented pathogenic microorganisms’ proliferation that usually characterized traditional way of its production. The behavioural coordination of pH and the total titratable acidity during fermentation reflected on the availability of the microbial metabolism to improve the fermentation rate. Furthermore, it was revealed that the colour measurements of some samples fermented with mutant strains of *Bacillus subtilis*, which is usually used to determine food acceptance were improved and better than the control samples.

Introducing a natural seasoning to serve as functional food in combating the side effects of consumption of artificial seasoning linked to harmful effects that has become a global cause of health issues, which necessitated this research, aside serving as a low-cost protein substitute.

## Introduction

Plants with oily-based seeds are high in protein and bioactive compounds with functional roles, which contributed to African diet meals (Polak et al., 2015). One of these oily-based seeds is *Citrullus vulgaris* seeds differently processed into various food condiments /seasonings such as *ogiri*. Food condiments fermented are usually characterized with strong objectionable odour, which do give sweetening taste to soups, sauces and other prepared dishes (Ogunshe et al., 2008). Condiments are usually serve as protein substitute and calorie intake in the diet to prevent malnutrition, especially in children (Nazidi et al., 2018). Several researchers have reported that *ogiri* are traditionally produce wild uncontrollable solid-state fermentation (Achi, 2005), known to have pathogenic and spoilage microbial contaminant with off-flavour (Ogueke and Nwagwu, 2007.

Fermentation play key roles in improving fermented product as a result of biochemical activities that occur which will bring about enhanced sensory attributes such as appearance, taste and aroma and improved nutritional properties (Sharma et al., 2020; Sanlier et al., 2019). Proteolytic *B. subtilis* has been identified as the major microorganism responsible for fermentation of *ogiri*. Microbial fermentation of *C. vulgaris* to produce *ogiri* can be natural or by the use of starter culture (s). Different microorganisms are usually isolated from the natural fermented or uncontrolled *ogiri* production with high microbial loads, but fewer fermenting microorganisms are isolated from the use of starter culture.

Mutation of microbial cells for food production has been in existence foe ages with various improved techniques for metabolic properties of starter culture selections. Various mutagenic agents such as UV irradiations and some chemicals are employed, which increased mutational rate from 10^1^ to 10^5^ cells per generation depending either for auxotrophic (primary) or for secondary mutants higher than natural change in chromosomal cell of 10^6^ to 10^7^ (Harlander 1992). There have been intense shortcomings with various agents to cause mutation such as time-consuming, imprecise, expensive and deleterious noticed mutation, etc. Notwithstanding the challenges, mutation has widely caused an improvement on the role of beneficial microorganisms, especially probiotic microorganisms in food production. Babatuyi et al. (2023) buttressed this point that using mutagenic agents produced mutant *B. subtilis* to release D-ribose to ameliorate inflammatory cytokines with increased ATP synthesis to sustain the tested animal (Wistar albino rats). It has also been reviewed by others researchers that D-ribose ameliorated kidney dysfunction and function as nucleotide flavour enhancers, adenine salvage rates in skeletal muscle recovery of intense contractions in both human and animal (Bayram et al., 2015; Kerksick et al., 2018; Li et al., 2021).

Colour analysis is a crucial parameter that differentiate between shades of colour among food products after treatment, where colour selection is important for consideration as one of the acceptance tools apart from the cost and quality. However, consumers are easily influenced by foreknowledge of how acceptable seasoning should appear and to improve upon on the natural production to serve as functional foods. The colour is measured by colorimeter to reflects the three parameter numbers (L*a* b*) known as CIELAB and the total colour change (ΔE) which is widely used in food industries and globally recognized. (Kortei et al., 2015; Hunterslab Technical Manual, 2008). The colorimetric parameter is widely used to characterize colour variation after processing conditions (Maskan, 2001).

*Ogiri* in its present form is only applied as flavour enhancer and its function can be enhanced as functional *ogiri* from the above statement if fermenting microorganism can be mutated in relation to the increase in quest for safe and natural seasonings. This necessitated this research to ferment *ogiri* with mutant and non-mutant strains of *Bacillus subtilis* from *Citrullus vulgaris* seeds and its effects on microbial population and colour characteristics of the fermented product.

## Materials and Methods

### Source of *Citrullus vulgaris*

Seeds of *Citrullus vulgaris* (‘Egusi’) authenticated by Crop, Soil and Pest Management Department, Federal University of Technology Akure were bought from the Owena market, Osun State, Nigeria. Already processed *ogiri* (market sample) were purchased from Oja-Oba in Akure, Ondo State, Nigeria. They were purchased in the month of August, 2020. Large quantity of the *C. vulgaris* seeds measured thirty kilograms (30 kg) were purchased and processed as a batch and used for the production of the *ogiri*.

### Reagents and Chemicals

The reagents and chemicals used were xylose, glucose, D-ribose HPLC standard, sodium chloride, 0.1 M potassium phosphate buffer (pH 7.0), sodium hydroxide (0.1 M Na0H), phenolphthalein indicator, Microbiological media [Tryptone Soy Agar (TSA), and Luria Broth (LB)] were OXOID. All were purchased from Sigma-Aldrich Chemie (Steinem, Germany) and of analytical grade. Distilled water was sterilized before use.

### Microbiological Analyses

#### Bacteria isolation

Strains of *B. subtilis* were isolated from the market *ogiri* sample by standard microbiological techniques (Fawole and Oso, 2007), using Tryptone Soy agar (OXOID), and the bacteria isolated were identified based on cultural, morphological and biochemical characteristics (Cowan, 2002).

#### Molecular characterization

The DNA extraction (16S rRNA gene analysis) and Polymerase chain reaction (PCR) of extracted DNA from isolated bacteria using universal primers 27F (5’-AGAGTTTGATCCTGGCTCAG-3’) and 1392R (5’-GGTTACCTTGTTACGACT-3’) were determined according to the methods of Akinyemi and Oyelakin (2014). The 16S rRNA sequences of both mutant and non-mutant strains were aligned using the Codon Code Aligner software (Codon Code Corporation, Centerville, MA) for mutation detection.

#### Standardization of starter culture

Identified bacteria and their standardization for starter culture were carried out as described by Babatuyi et al. (2020) using McFarland standard. McFarland Standard turbidity was visually compared to bacterial suspension of cultured broth and the approximate concentration of the bacteria in suspension. The turbidity of each bacterial suspension was measured with spectrophotometer at 600 nm (model DNP-9102 721-VIS Series SEARCHTECH instruments, England). The equation is shown below:

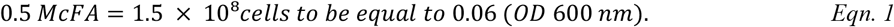

#### Preparation of starter culture

The starter culture was prepared as described by Babatuyi et al. (2020) with modification. Eighteen (18) h old broth culture of one milliliter (1 mL) of cell suspension (approximately 1.5 × 10^8^cfu/mL) harvested was washed three times by suspending in 10 ml 0.1 M potassium phosphate buffer (pH 7.0) and centrifuged using refrigerated centrifuge (Model Harrier 18/80, Henderson Biomedical LTD MSE, UK) at 5000 g for 10 min. Re-suspension of the cells in 20 mL sterile 0.1M potassium phosphate buffer (pH 7.0) were aseptically carried out and standardized using a spectrophotometer (Model V-721 VIS, China) at 600 nm wavelength approximately *1*.*5 × 10*^*8*^ of 1.0 mL of the cell suspension.

### Production of mutant strains of *Bacillus subtilis*

#### Physical mutation

Mutant strains of *B. subtilis* were produced by using the modified method of Babatuyi et al. (2023). Four milliliters (4 mL) of cell suspension (10^8^ cells/mL) of *B. subtilis* was placed in Petri dish under ultraviolet (UV) lamps (253.7nm and 366nm) at a distance of 30 cm and irradiated at time of 10 s intervals from 30 s to 120 s. Each cell suspension was shaken during irradiation. The cells were diluted in sterile physiological saline (0.85 %) and 1 mL of each cell suspension was plated on Luria Broth, incubated using electro-thermal incubator (Model DNP SEARCHTECH instruments, England) in the dark at 37 ° C for 2 days.

#### Chemical mutation

Mutant strains of *B. subtilis* were produced using chemical mutagen by the modified method of Babatuyi et al. (2023). One milliliter (1 mL) of SDS solution (100 mg/mL) was added to 1 mL of the cell suspension of each *B. subtilis* strain (10^8^ cells/mL). It was placed in rotary shaker at time of 10 s intervals from 30 to 120 s. The mixture was diluted several times with sterile distilled water immediately to terminate the reaction. The cells were diluted in sterile physiological saline. The plates were incubated at 37 ° C in dark for 3 days.

#### D-ribose production

Five milliliter (5 mL) of seeded microbial broth culture of mutant and non-mutant media containing 5 g/L xylose, 5 g/L glucose each was prepared and incubated for 48 h at 37 ° C in water bath shaker (constant temperature oscillator) model HA-C2 Labscience, England). The sugar composition of the culture broth was analyzed after incubation using HPLC-RI according to the method of Park and Seo (2004).

#### Production of *ogiri*

The *ogiri* sample was produced as described by Akinyele and Oloruntoba (2013) with modification. *C. vulgaris* seeds purchased from the market were sorted and washed. The seeds were boiled for 40 min to remove the outer coat and drained. The de-hulled seeds were re-boiled for another 2 h to soften the seeds. The seeds were mashed and divided into 9 portions. Eight (8) out of the nine (9) portions were fermented with mutant strains of *B. subtilis*, while the last portion was fermented with non-mutant strain to serve as the control and as well as the market sample. The samples were wrapped in different low-density polyethylene packaging (LDP) materials, before they were fermented at 28 ^o^ C for 5 days. After fermentation, the samples were re-wrapped in low-density polyethylene packaging (LDP) materials. They were cured by placing them in a warm dry enclosure for 21 days. The cured *ogiri* samples were then freeze-dried (Lab Kit FD-10-MR), milled, packaged and stored for further analysis.

#### Chemical Analyses

##### pH

The pH of *ogiri* fermented with mixed starter cultures was determined using Kent pH meter (Kent Ind. Measurement Ltd., UK) model 7020 with a glass electrode according to the method of A.O.A.C. (2015) and calculated as lactic acid.

##### Total Titratable Acidity (TTA)

The total titratable acidity of *ogiri* fermented with mutant and non-mutant starter cultures was determined according to the method of A.O.A.C. (2015).

The mean of TTA obtained from triplicate determination was calculated as follows:

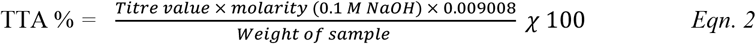

##### Colour Measurements

The characteristic of the colour was determined by McGuire (1992) using colorimeter (model PCE-CSM2, American). The CIE L*a*b* colour coordinates degree of lightness (L*) ranged from zero (black) to 100 (white), red/green (a*), ranged from +60 (red) to -60 (green), yellow/blue (b*) ranged from +60 (yellow) to –60 (blue). The Chroma (C) estimates colour saturation or intensity, while the hue angle (Ho) reports the relative amounts of redness and yellowness where 0 °/360 ° is defined for red/magenta, 90 ° for yellow, 180 ° for green and 270° for blue colour or purple, or intermediate colours between adjacent pairs of these basic colours. The colour difference (ΔE) was calculated in relation to the control sample (Saricoban and Yilmaz, 2010) as follows:

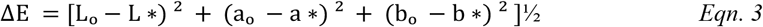

where L_0_, a_0_ and b_0_ are values for initial, while L*, a* and b* are values for the final sample.

### Data analysis

The results of the *ogiri* samples analyzed were pooled and expressed as mean ± standard deviation (SD). Mean values were analyzed and compared using one-way ANOVA followed by New Duncan Multiple Range Test (NDMRT). The significance was at p ≤ 0.05.

## Results and Discussion

### Cultural, morphological and biochemical characteristics of bacterial isolated during ogiri fermented with mutant and non-mutant strains of *Bacillus subtilis* and market sample from *Citrullus vulgaris* seeds

A total of 12 different genera of microorganisms were isolated from the *ogiri* samples which includes: *Bacillus subtilis, Bacillus cereus, Bacillus pumilus, Bacillus megatarium, Bacillus licheniformis, Enterococcus faecalis, Staphylococcus aureus, Lactobacillus* sp. *Proteus* sp. and enterobacteriaecea (*Salmonella* sp., *Shigella* sp. and *Escherichia coli*). The cultural, morphological and biochemical characteristics are shown in Table 1.

**Table 1:**
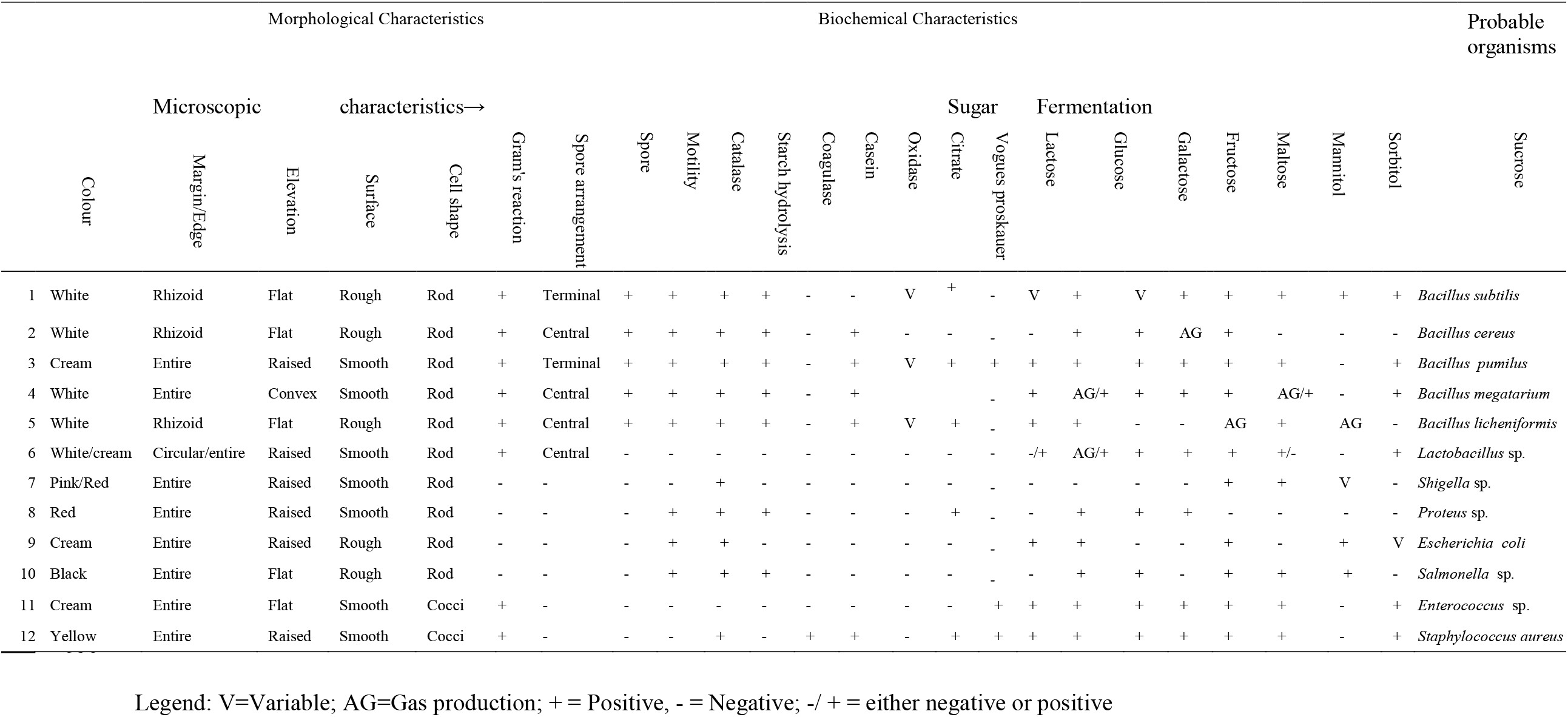
Cultural, morphological and biochemical characteristics of bacterial isolated from ‘ogiri’ during *Citrullus vulgaris* seeds fermentation with mutant and non-mutant strains of *Bacillus subtilis*

### Molecular identification of mutant and non-mutant strains of *Bacillus subtilis* used for the production of *ogiri* from *Citrullus vulgaris* seeds

The molecular identification of Mutant and non-mutant Stains of *Bacillus subtilis* used to produce *ogiri* from *Citrullus vulgaris* is shown in Table 2. It was observed that after molecular identification of eight (8) *Bacillus subtilis* with higher D-ribose production after mutation and non-mutant strain, five (5) strains were identified. Three (3) mutant strains had the same strains with the parent *Bacillus subtilis* (NMS00), while two from physical mutant had the same strains.

**Table 2:** Molecular identification of mutant and non-mutant strains of *Bacillus subtilis* used for the production of *ogiri* from *Citrullus vulgaris* seeds.

| S/N | *Bacillus subtilis* Code | MUTAGENIC AGENT | MOLECULAR IDENTITY |
| --- | --- | --- | --- |
| 1 | MIC67 | Physical mutation | *B. subtilis* EBS/SRLAAH/2017 |
| 2 | MIC64 | Physical mutation | *B. subtilis* strain Saad27 |
| 3 | MIC21 | Physical mutation | *B. subtilis* 16 S KH28.1 |
| 4 | MHL23 | Physical mutation | *B. subtilis* strain SE10C10 |
| 5 | MHL22 | Physical mutation | *B. subtilis* strain Saad27 |
| 6 | MSD99 | Chemical mutation | *B. subtilis* strain SE10C10 |
| 7 | MSD51 | Chemical mutation | *B. subtilis* strain P2 |
| 8 | MSD30 | Chemical mutation | *B. subtilis* strain SE10C10 |
| 9 | NMS00 | Control | *B. subtilis* strain SE10C10 |

MIC67: *Bacillus subtilis* after mutation to give *B. subtilis* EBS/SRLAAH/2017 strain; MIC64: *Bacillus subtilis* after mutation to give *B. subtilis* Saad27 strain; MIC21: *Bacillus subtilis* after mutation to give *B. subtilis* S KH28.1 strain; MHL23 *Bacillus subtilis* after mutation to give *B. subtilis* strain SE10C10 strain. MHL22: *Bacillus subtilis* after mutation to give *B. subtilis* strain Saad27 strain; MSD99: *Bacillus subtilis* after mutation to give *B. subtilis* strain SE10C10 strain; MSD51: *Bacillus subtilis* after mutation to give *B. subtilis* strain P2 strain; MSD30: *Bacillus subtilis* after mutation to give *B. subtilis* strain SE10C10 strain; NMS00: *Bacillus subtilis* without mutation to give *B. subtilis* strain SE10C10 strain

### Microbial Load of O*giri* Samples Fermented with Mutant and Non-mutant Strains of *Bacillus subtilis* with Market Sample during Fermentation

The total microbial load of the *ogiri* samples fermented with mutant and non-mutant strains of *Bacillus subtilis* are shown in Figure 1. The total microbial load at day 1 ranged from 1.0 × 10^3^ to 3.0 × 10^4^ cfu/g. There was a steady increase from day 2 (1.0 ×10^4^ to 3.0 ×10^4^ cfu/g) to day3 (1. 4 ×10^4^ to 3.6 ×10^4^ cfu/g) before decreasing from day 3 (1.3 ×10^4^ to 3.9 ×10^4^ cfu/g) to the last day of fermentation (1.1 ×10^3^ to 1.0 ×10^4^ cfu/g. This could be due to the microbial activities during growth and the increase in TTA (fermentation indictor) towards the end of the fermentation. The growth’ pattern moved from lag phase, where they are metabolically active and acclimatizing to the environment before adaptation through exponential cell divisions. The population growth got to the stationary phase at the end of the fermentation on day 3 and decreased as it enters death phase on day 4 due release and accumulation of toxic wastes (Babatuyi et al., 2019), dead cells and depletion of the substrate to the end of fermentation (Vermeersch, et al., 2019; Liu et al, 2016).

**Figure 1.**
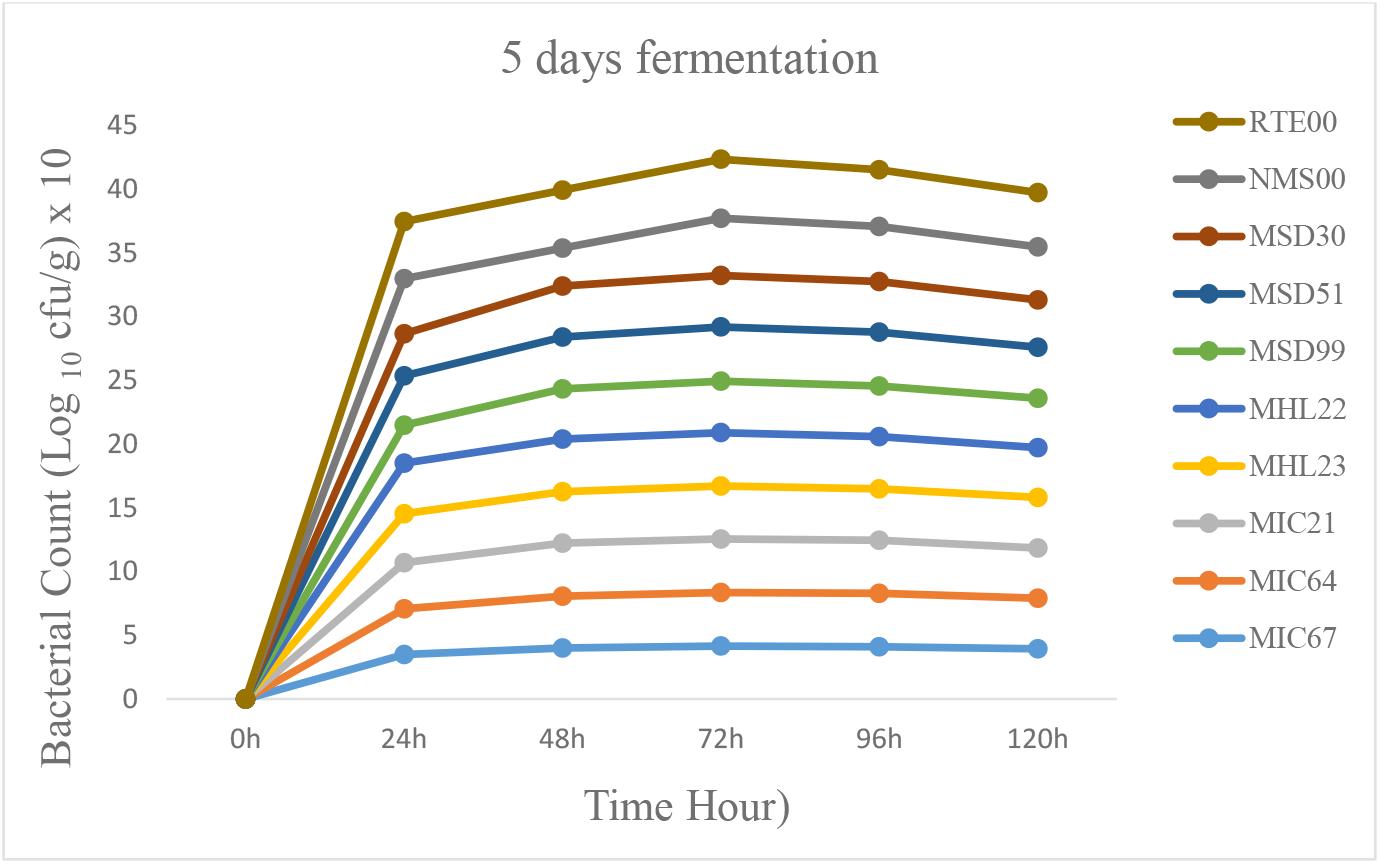
Microbial load of ‘ogiri’ during fermentation of *Citrullus vulgaris* seeds with mutant and non-mutant strains of *Bacillus subtilis*. Legend: MIC67: *Ogiri* produced with *Bacillus subtilis* mutant strain exposed to inoculating chamber UV light at 90 sec, MIC64: *Ogiri* produced with *Bacillus subtilis* mutant strain exposed to inoculating chamber UV at 30 sec, MIC21: *Ogiri* produced with *Bacillus subtilis* mutant strain exposed to inoculating chamber UV at 120 sec, MHL23: *Ogiri* produced with *Bacillus subtilis* mutant strain exposed to hand UV lamp at 100 sec, MHL22: *Ogiri* produced with *Bacillus subtilis* mutant strain exposed to hand UV lamp at 110 sec, MSD99: *Ogiri* produced with *Bacillus subtilis* mutant strain exposed to SDS at 100 sec, MSD51: *Ogiri* produced with *Bacillus subtilis* mutant strain exposed to SDS at 110 sec, MSD30: *Ogiri* produced with *Bacillus subtilis* mutant strain exposed to SDS at 120 sec, NMS00: *Ogiri* produced with Non-mutant *Bacillus subtilis* strain without D-ribose yield, and RTE00: market sample.

### Microbial Occurrence of Laboratory Produced Ogiri Fermented with Mutant, Non-mutant Strains of *Bacillus subtilis* and Market Sample from *Citrullus vulgaris* Seeds

The occurrence of microorganisms isolated from the *ogiri* samples fermented with mutant and non-mutant strains with different media (agars) used are shown in Table 3. These populations characterized only bacteria species without fungi, which is similar to the report of Falegan (2011) who worked on *ogiri* produced in different states of southwest, Nigeria with *C. vulgaris*.

**Table 3:**
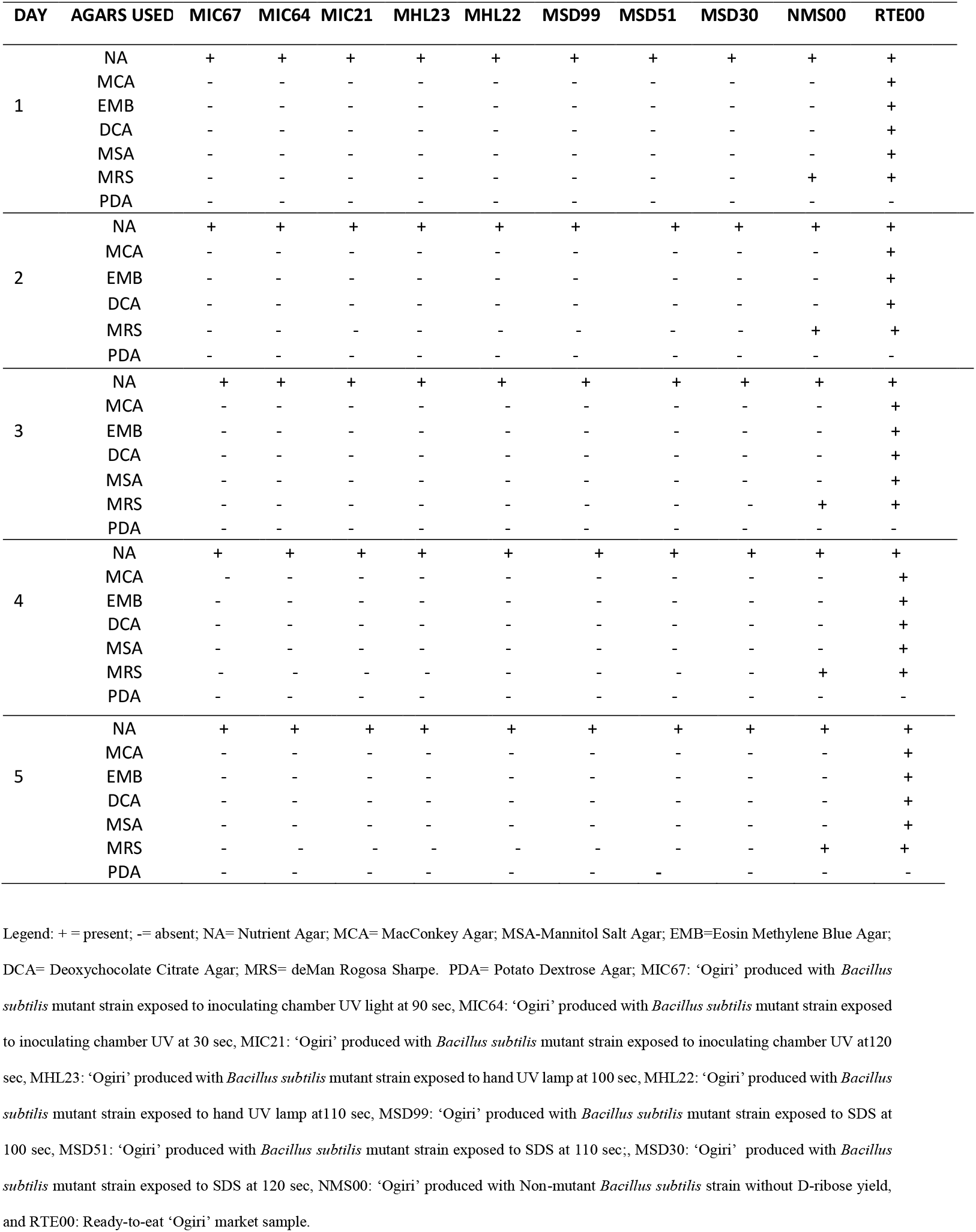
Occurrence of microorganisms in ‘ogiri’ produced by the fermentation of *Citrullus vulgaris* seeds fermentation with mutant and non-mutant strains of *Bacillus subtilis*.

During fermentation, species of *Bacillus* and few enterobacteriaecea (sample RTE) were found and as the fermentation progress, so also the microorganisms’ increased. The microbial loads were minimum compared to sample RTE00 (market sample), which was dominated by enterobacteriaecea. *Bacillus subtilis* dominated *ogiri* samples fermented with mutant *Bacillus subtilis* strains throughout the fermentation. This could be as a result of metabolic activities of mutant strains used for the fermentation which did not allow the growth of any other microorganism. Contamination is being introduced by the food processors during food handling and under poor hygiene (Adedeji *et al*., 2017).

### pH and Total Titratable Acidity (TTA) of O*giri* Samples Fermented with Mutant and Non-mutant Strains of *Bacillus subtilis* with Market Sample

The pH of the *ogiri* samples fermented with mutant and non-mutant strains of *Bacillus subtilis* is shown in Figure 2. The pH of the ogiri samples increased from 4.08 to 9.86 as the days of fermentation progressed to the last day of fermentation. The pH of the condiments aligned with works of Adesanya et al. (2021) on locus beans fermented with *B. subtilis* broth at both 1 ml (6.68 to 9.23) and 1.5 ml (6.94 to 8.98) concentrations. Ibeabuchi et al. (2014) work on both locus beans (6.31 to 7.20) and castor oil bean (6.36 to 7.15), and as well as Ogueke et al. (2013), on melon seeds (5.2 to 7.81) aligned. The involvement of *B. subtilis* contributed to the release of ammonia which cause objectionable odour due to breakdown of protein to amino acids, peptides and free ammonia (Achi, 2005) with increased proteinase and deaminase production Ogueke et al., 2013). This fasten the fermentation rate to result into well-fermented products.

**Figure 2.**
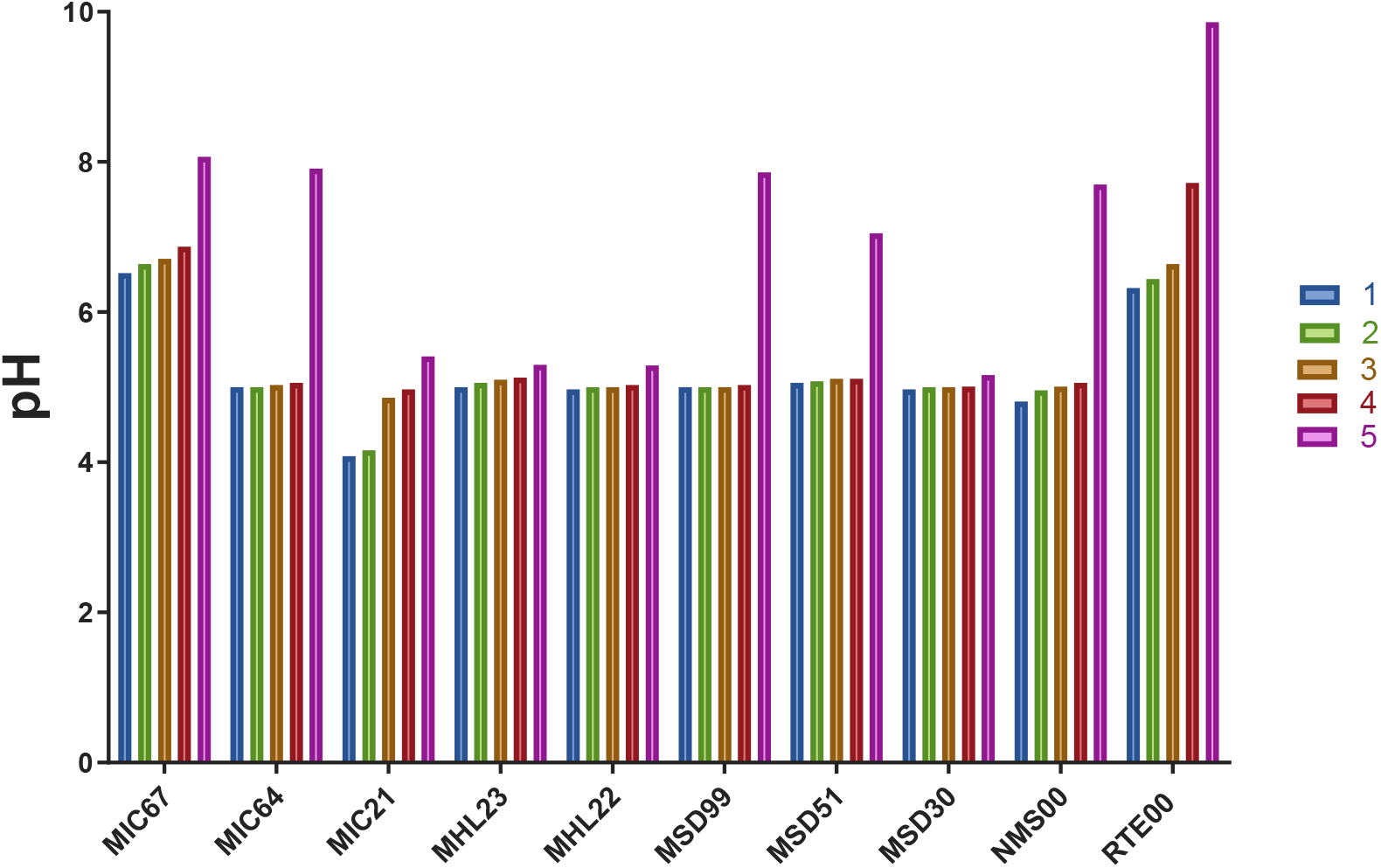
Changes in pH of *ogiri* samples during fermentation of *Citrullus vulgaris* seeds with mutant and non-mutant strains of *Bacillus subtilis*. Legend: MIC67: *Ogiri* produced with *Bacillus subtilis* mutant strain exposed to inoculating chamber UV light at 90 sec, MIC64: *Ogiri* produced with *Bacillus subtilis* mutant strain exposed to inoculating chamber UV at 30 sec, MIC21: *Ogiri* produced with *Bacillus subtilis* mutant strain exposed to inoculating chamber UV at 120 sec, MHL23: *Ogiri* produced with *Bacillus subtilis* mutant strain exposed to hand UV lamp at 100 sec, MHL22: *Ogiri* produced with *Bacillus subtilis* mutant strain exposed to hand UV lamp at 110 sec, MSD99: *Ogiri* produced with *Bacillus subtilis* mutant strain exposed to SDS at 100 sec, MSD51: *Ogiri* produced with *Bacillus subtilis* mutant strain exposed to SDS at 110 sec, MSD30: *Ogiri* produced with *Bacillus subtilis* mutant strain exposed to SDS at 120 sec, NMS00: *Ogiri* produced with Non-mutant *Bacillus subtilis* strain without D-ribose yield, and RTE00: market sample.

The trend at which pH of sample RTE00 increased throughout the fermentation could be as a result of many microorganisms breaking down the substrate simultaneously during fermentation to give more objectionable odour compared to samples fermented with mutant and non-mutant strains with steady increase. However, sample MIC67 had the same trend, making it closer to sample RTE00.

The TTA of the *ogiri* samples fermented with mutant and non-mutant strains of *Bacillus subtilis* is shown in Figure 3. The TTA does not follow the same pattern in all the samples. Samples MIC67 (0.40 to 1.30 %); MIC64 (0.40 to 1.50 %); MSD99 (0.30 to 1.50 %); MSD51 (0.40 to 1.50 %) and NMS00 (0.40 to 1.50 %) have their pH (6.53 to 8.67; 5.00 to 7.91; 5.00 to 7.86; 5.06 to 7.05 and 4.81 to 7.70) increased throughout the fermentation respectively.

**Figure 3.**
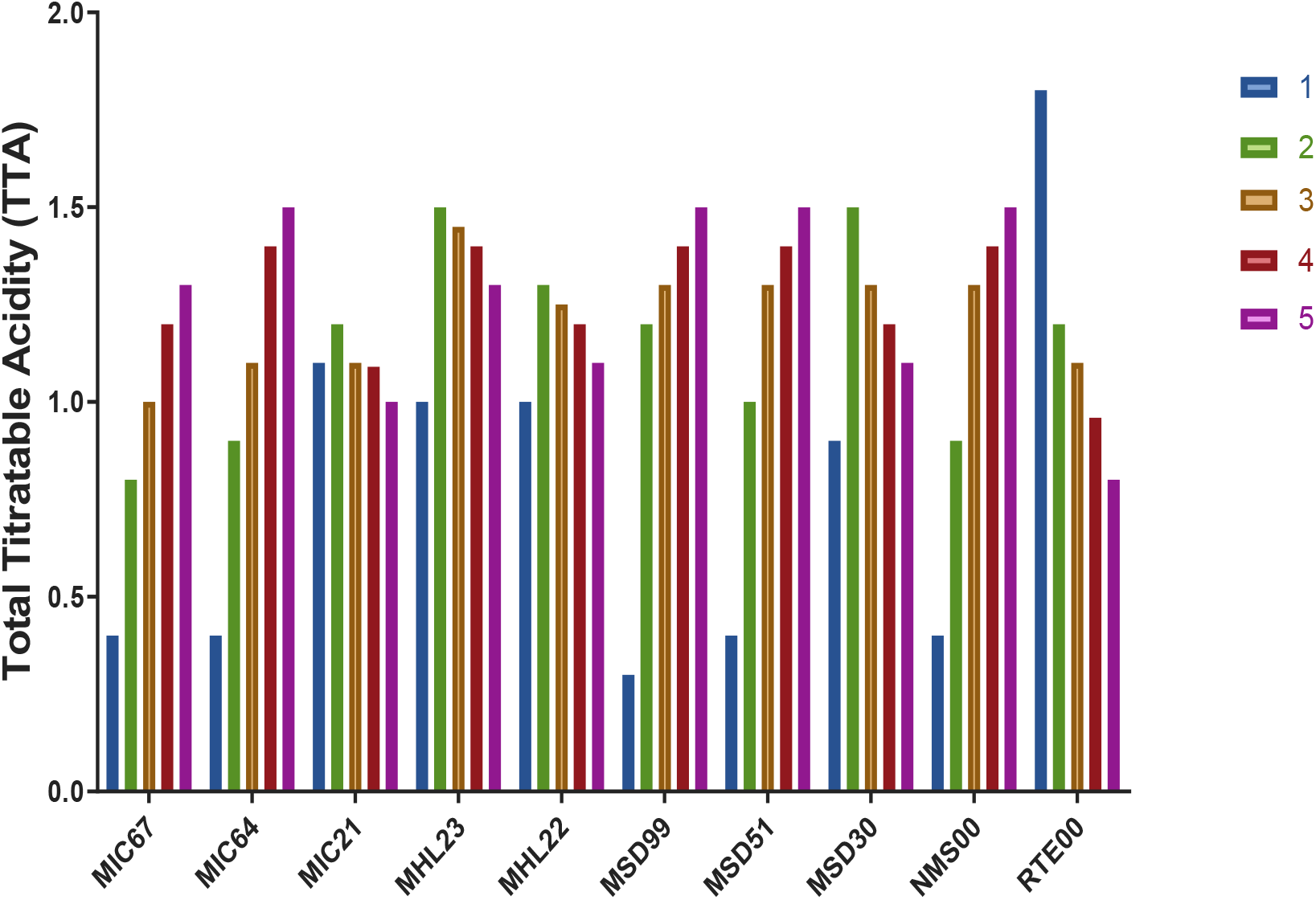
Changes in Total Titratable Acidity (TTA) of *ogiri* samples during fermentation of *Citrullus vulgaris* seeds with mutant and non-mutant Strains of *Bacillus*. Legend: MIC67: *Ogiri* produced with *Bacillus subtilis* mutant strain exposed to inoculating chamber UV light at 90 sec, MIC64: *Ogiri* produced with *Bacillus subtilis* mutant strain exposed to inoculating chamber UV at 30 sec, MIC21: *Ogiri* produced with *Bacillus subtilis* mutant strain exposed to inoculating chamber UV at 120 sec, MHL23: *Ogiri* produced with *Bacillus subtilis* mutant strain exposed to hand UV lamp at 100 sec, MHL22: *Ogiri* produced with *Bacillus subtilis* mutant strain exposed to hand UV lamp at 110 sec, MSD99: *Ogiri* produced with *Bacillus subtilis* mutant strain exposed to SDS at 100 sec, MSD51: *Ogiri* produced with *Bacillus subtilis* mutant strain exposed to SDS at 110 sec, MSD30: *Ogiri* produced with *Bacillus subtilis* mutant strain exposed to SDS at 120 sec, NMS00: *Ogiri* produced with Non-mutant *Bacillus subtilis* strain without D-ribose yield, and RTE00: market sample.

This could be due to both genetic and physiological resistance that have transformed and changed the microorganisms responsible for the fermentation to survive in acidic environments that prevailed during the fermentation (Guan and Liu, 2020). The TTA of samples MIC21, MHL23, MHL22 and NMS00 increased to day 2 of fermentation before decreasing towards the end of fermentation. The results could due to microbial activities changing the fermentation pattern as TTA is an indicator for proper fermentation of the substrate.

### Colour Measurements

The results of colour measurements of *ogiri* samples after production is presented in Table 4. The whiteness value (L*) ranged from 18.17±0.00^j^ to 51.93±0.00^a^, red/ green value (a*) ranged from 3.19 to11.49±0.00 and yellow /blue (b*) ranged from 4.56 to 25.55±0.00 respectively. There was significant (P<0.05) difference in all the parameters of L*, a* and b* with respect to different mutant agents and exposure time. The o*giri* samples were higher in these parameters than the control sample (RTE00): 18.17±0.00^j^, 3.19±0.00^j^ and 4.56±0.00^j^ respectively. This is not about the colour characterization, but also the arrangement of colour space in the quadrant coordinate. The lower L* value of control sample (18.17±0.00^j^) could be due to the reaction of Maillard between proteins and reducing sugars. This was due to the microbial activities during fermentation and curing which imparted the brownish black colour (Shaf et al. 2016). The pattern of increase is similar to results of Koitei et al. (2015) on determination of colour parameters of gamma irradiated fresh and dried mushrooms during storage (60.46 to 61.30; 3.47 to 3.91 and 18.13 to 19.39). Jan et al. (2022) also buttressed the statement with similarities in their research on influence of replacement of wheat flour by rice flour on rheo-structural changes, *in vitro* starch digestibility and consumer acceptability of low-gluten pretzels, having (43.57±2.12 to 56.49±2.37; 8.71 ± 1.67 to 9.43 ± 1.73 and 22.33 ± 1.24 to 17.59 ± 1.19) respectively.

**Table 4:** Colour Measurements of *Ogiri* Samples during Fermentation of *Citrullus vulgaris* seeds with Mutant and Non-mutant Strains of *Bacillus subtilis*.

|  | L* | a* | b* | c* | h* | ΔE |
| --- | --- | --- | --- | --- | --- | --- |
| MIC67 | 21.84 | 9.48 | 13.19 | 16.25 | 54.31 | 11.29 |
| MIC64 | 38.55 | 10.38 | 20.12 | 22.64 | 62.70 | 26.62 |
| MIC21 | 51.40 | 10.62 | 25.44 | 27.57 | 67.33 | 39.94 |
| MHL23 | 35.27 | 9.34 | 21.61 | 23.55 | 66.62 | 24.92 |
| MHL22 | 45.25 | 10.77 | 25.55 | 27.72 | 67.13 | 35.09 |
| MSD99 | 33.36 | 9.22 | 19.48 | 21.55 | 64.67 | 22.13 |
| MSD51 | 35.11 | 11.49 | 21.19 | 25.88 | 63.64 | 25.15 |
| MSD30 | 50.93 | 9.41 | 23.15 | 24.99 | 67.88 | 38.18 |
| NMS00 | 31.16 | 10.34 | 19.66 | 22.21 | 62.25 | 21.16 |
| RTE00 | 18.17 | 3.19 | 4.56 | 5.56 | 55.04 | 0.00 |

Legend: L\* = Degree of lightness; a\* = Degree of redness/purple hue; b\* = Degree of yellowness/bluish hue; C\* = Chroma; h° = hue angle; ΔE = Total colour change; MIC67: *Ogiri* produced with *Bacillus subtilis* mutant strain exposed to inoculating chamber UV light at 90 sec, MIC64: *Ogiri* produced with *Bacillus subtilis* mutant strain exposed to inoculating chamber UV at 30 sec, MIC21: *Ogiri* produced with *Bacillus subtilis* mutant strain exposed to inoculating chamber UV at 120 sec, MHL23: *Ogiri* produced with *Bacillus subtilis* mutant strain exposed to hand UV lamp at 100 sec, MHL22: *Ogiri* produced with *Bacillus subtilis* mutant strain exposed to hand UV lamp at 110 sec, MSD99: *Ogiri* produced with *Bacillus subtilis* mutant strain exposed to SDS at 100 sec, MSD51: *Ogiri* produced with *Bacillus subtilis* mutant strain exposed to SDS at 110 sec, MSD30: *Ogiri* produced with *Bacillus subtilis* mutant strain exposed to SDS at 120 sec, NMS00: *Ogiri* produced with Non-mutant *Bacillus subtilis* strain without D-ribose yield, and RTE00: market sample.

The Chroma (C), hue angle (H) and total colour change ranged from 5.56±0.00^j^ to 27.720.00^a^, 54.31±0.00^j^ to 67.88±0.00^j^ and 11.29±0.00^i^ to 39.94±0.00 respectively. The o*giri* samples with higher ΔE implies greater colour change in food products, hence acceptable to consumers. Difference in the low total colour change (ΔE) values of the *ogiri* samples could be lack of adequate water activity needed for enzymatic metabolic activities to form melanin pigmented dark colour (Shafi et al. 2016).

## Conclusion

The effects of production of *ogiri* fermented with mutant and non-mutant of *Bacillus subtilis* and Market samples on microbial population and colour characteristics has been reported. It was observed that *B. subtilis* predominates in *ogiri* fermented with mutant strain of *B. subtilis* with no spoilage and pathogenic which was inherent in the market sample of *ogiri* that was naturally fermented. These were reflected in the behaviour of their chemical composition, which could be as a result of D-ribose production. These *ogiri* samples had Lightness (L*) higher than the control, an indicator for acceptability. Integration of these modified *ogiri* samples into the food system with improved quality could enhance promoting large scale industrialization.

## Abbreviations

UV: Ultraviolet irradiation
SDS: Sodium Dodecyl Sulphate
MIC67: *Ogiri* produced with *Bacillus subtilis* mutant strain exposed to inoculating chamber UV light at 90 sec
MIC64: *Ogiri* produced with *Bacillus subtilis* mutant strain exposed to inoculating chamber UV at 30 sec
MIC21: *Ogiri* produced with *Bacillus subtilis* mutant strain exposed to inoculating chamber UV at 120 sec
MHL23: *Ogiri* produced with *Bacillus subtilis* mutant strain exposed to hand UV lamp at 100 sec
MHL22: *Ogiri* produced with *Bacillus subtilis* mutant strain exposed to hand UV lamp at 110 sec
MSD99: *Ogiri* produced with *Bacillus subtilis* mutant strain exposed to SDS at 100 sec
MSD51: *Ogiri* produced with *Bacillus subtilis* mutant strain exposed to SDS at 110 sec
MSD30: *Ogiri* produced with *Bacillus subtilis* mutant strain exposed to SDS at 120 sec
NMS00: *Ogiri* produced with Non-mutant *Bacillus subtilis* strain without D-ribose yield
RTE00: Market sample

## Acknowledgements

The authors are grateful to the Departments of Microbiology and Food Science and Technology of Federal University of Technology in Akure, Nigeria for providing additional equipment and facilities.

## Author’s contributions

CYB analyzed and interpreted the data, acquisition, design the work, drafted the work, wrote the manuscript; VOO acquisition, substantively revised it; FAA acquisition, substantively revised it. We all agreed to be accountable for the author’s own contributions and to ensure that questions related to the accuracy or integrity of any part of the work, even ones in which the author was not personally involved, are appropriately investigate, resolved, and the resolution documented in the literature. All authors approved the final version of the manuscript.

## Availability of data and materials

The datasets used and/or analyzed during the current study are available from the corresponding author on reasonable request

## Declarations

